# Leucine rich repeat kinase 2 (LRRK2) gly2019ser mutation is absent in a second cohort of nigerian africans with parkinson disease

**DOI:** 10.1101/363945

**Authors:** Njideka U Okubadejo, Mie Rizig, Oluwadamilola O Ojo, Hallgeir Jonvik, Olajumoke Oshinaike, Emmeline Brown, Henry Houlden

## Abstract

To date the LRRK2 p.G2019S mutation remains the most common genetic cause of Parkinson disease (PD) worldwide. It accounts for up to 6% of familial and approximately 1.5% of sporadic cases. LRRK2 has a kinase enzymatic domain which provides an attractive potential target for drug therapies and LRRK2 kinase inhibitors are in development. Prevalence of the p.G2019S has a variable ethnic and geographic distribution, the highest was reported among Ashkenazi Jews (30% in patients with familial PD, 14% in sporadic PD, 2.0% in controls) and North African Berbers (37% in patients with familial PD, 41% in sporadic PD, and 1% in controls). Little is known about the frequency of the LRRK2 p.G2019S among populations in Sub Saharan Africa. Our group and others previously reported that the p.G2019S is absent in a small cohort from Nigerian PD patients and controls. Here we used Kompetitive Allele Specific PCR (KASP) assay to screen for the p.G2019S in a larger cohort of Black African PD patients (n =126) and healthy controls (n = 55) from Nigeria. Our analysis confirmed that all patients and controls are negative for the p.G2019S mutation. This report provides further evidence that the LRRK2 p.G2019S is not implicated in PD in black populations from Nigeria and support the notion that p.G2019S mutation originated after the early human dispersal from sub-Saharan Africa. Further studies using larger cohorts and advance sequencing technology are required to underpin the genetic causes of PD in this region.

## INTRODUCTION

Mutations in the leucine-rich repeat kinase 2 (*LRRK2*) gene, initially described in 2004, are the most prevalent genetic cause of Parkinson disease (PD), and have been associated with both familial and sporadic cases of PD in some populations.(1,2) Although at least seven distinct pathogenic mutations in *LRRK2* have been proven to cause PD, there is considerable population variability in the frequencies of these mutations.(3) The most common is the LRRK2 p.G2019S substitution, with a global frequency of 1% in sporadic PD and up to 5% in familial PD.(4) The highest frequencies of LRRK2 p.G2019S mutations are reported from the North African Berber and Arab populations, where it is present in 30-42% of familial PD, 30-39% of apparently sporadic PD, and 0-2.2% of healthy controls.(5)

The interest in exploring genetic underpinnings to PD extend beyond elucidating pathogenetic mechanisms, to ultimately manipulating the knowledge to derive viable disease modifying and therapeutic targets. Following evidence of gain-of-function increase in LRRK2 kinase activity in mutation carriers, promising pre-clinical studies have already demonstrated neuroprotective properties of LRRK2 kinase inhibitors, including reduced alpha-synuclein aggregation, reduced dopaminergic neurodegeneration, and reduced microglial and macrophage activation.(6) Considering the enormous potential future application to LRRK2 p.G2019S –related PD, determining the relevance in diverse populations is imperative.

Although comprehensively studied in numerous European and North American populations, studies on the frequency of LRRK2 p.G2019S in sub-Saharan Africa have been scarce. A previous report from our Nigerian African population including 57 sporadic PD patients and 51 healthy controls did not identify LRRK2 p.G2019S mutations in either PD cases or controls. (7) In this study we aimed at exploring a larger cohort of PD cases and healthy controls from a Black African Nigerian background for the LRRK2 p.G2019S mutation to further clarify the role of this mutation in PD in sub Saharan Africans.

## METHODOLOGY

### Participant recruitment, diagnostic ascertainment and clinical evaluation

PD cases (n=126) were recruited from the Movement Disorders Clinic of the Lagos University Teaching Hospital, Idi Araba, Lagos State, Nigeria in period between February 2016 to May 2017. The International Parkinson Disease and Movement Disorder Society (IPMDS) clinical diagnostic criteria were applied in this study, and cases meeting the diagnostic criteria for both clinically established PD and probable PD were included.(8) Hoehn and Yahr stage was as per the IPMDS revised Unified Parkinson Disease Rating Scale.(9) Case ascertainment was rigorous and based on agreement by two movement disorders specialists. Both sporadic and familial PD cases were included. Persons with a presumed secondary cause or red flags suggestive of other hereditary of degenerative Parkinsonism were excluded from the study. The protocol included information on demographic characteristics, age at onset of initial motor symptom of PD, and family history of tremor or parkinsonism in a first or second degree relative. Age at onset was defined by the age (years) at onset of the earliest motor symptom of PD. Controls (n=55) were population-based unrelated volunteers recruited via advertisement to the neighbouring communities, and were age ± 3 years, gender-matched, otherwise neurologically healthy, and without any clinical evidence of medical illness (as determined by history and a medical and neurological clinical examination).

This study was approved by the Health Research Ethics Committee of the Lagos University Teaching Hospital, Idi Araba, Lagos (Ethics approval # ADM/DCST/HREC/366) and University College London and the National Hospital for Neurology and Neurosurgery-London (Ethics approval # 07/Q0512/26). In accordance with the protocol, written informed consent was obtained from all participants, and included permission to re-contact for future studies. Clinical data from hard-copy case record forms were logged into a secure database. All data was de-identified to protect confidentiality, and assigned a unique study code and given a unique laboratory code. Clinical variables were analysed using the IBM ^®^ Statistical Package for Social Sciences (SPSS) ^®^ version 21. Chi-square test and analysis of variance test were used to compare categorical parameters and age respectively between PD cases and controls. Level of significance was set at a p value <0.05.

### DNA extraction and LRRK2 p.G2019S genotyping

Whole blood (20 mls) was collected into EDTA Vacutainer bottles using standard protocols. Samples were stored at minus 80^0^C (degrees centigrade) in the Central Research Laboratory of the College of Medicine, University of Lagos, until sample shipment to the designated extraction laboratory. Genomic DNA was extracted using the standard diagnostic laboratory protocols. DNA concentration was measured with a NanoDrop Spectrophotometer (Thermo Scientific, Waltham, MA, USA). Samples were diluted to a uniform concentration, and the equivalent of 50 ng of DNA per sample were arranged in 96-well plates. Two negative controls (DNA empty wells) and two positive controls (a well containing DNA from a previously genotyped normal individual and a second well containing DNA from an individual who is known heterozygous for the p.G2019S mutation) were included in each plate. No positive homozygous control for the p.G2019S was available. The SNP of interest p.G2019S (rs34637584) and a 50 base-pairs (bp) sequence surrounding the SNP were annotated using ENSEMBL genome browser and were submitted to LGC Genomics website via their SNP submission template to design primers. Genotyping was performed using Kompetitive Allele Specific PCR genotyping assay (KASP™, LGC Genomics. Herts, UK) according to the manufacturer’s protocol. Genotyping results were visualized with SNP Viewer software (version 1.99, Hoddesdon, UK). Genotypes were examined for deviation from Hardy-Weinberg equilibrium using Chi-square test.

## RESULTS

### Baseline clinical characteristics of PD cases and controls

The baseline characteristics of the PD cases and controls are shown in Table 1. The study included 126 PD cases and 54 controls with comparable mean ages (years) at the time of the study (61.9 ± 9.9 and 64.6 ± 8.7 respectively) (ANOVA; p=0.14). The median age at onset of PD was 59.0 years, while median duration of PD was 48 months. A family history of parkinsonism or tremor in a first degree relative was present in 20 (15.9%) of the PD and none of the controls (as this was an exclusion criterion for controls).

**Table 1.** Clinical characteristics of Parkinson disease cohort

| Characteristic | Parkinson<br>disease<br>n = 126 | Controls<br>n = 54 | Statistics<br>(p value) |
| --- | --- | --- | --- |
| Gender distribution, *n* (%) |  |  |  |
| *Male* | 93 (73.8) | 34 (63.0) | 0.14 |
| *Female* | 33 (26.2) | 20 (37.0) |  |
| Age at study (range), *years* | 36 – 81 | 41 – 80 |  |
| Age at study (mean ± SD), *years* | 61.9 ± 9.9 | 64.6 ± 8.7 | 0.09 |
| Age at study median), *years* | 63.5 | 64.0 |  |
| Age at onset (range), *years* | 33 – 80 |  |  |
| Age at onset (mean ± SD), *years* | 57.1 ± 9.9 |  |  |
| Age at onset (median), *years* | 59.0 |  |  |
| Young onset (≤ 50 years at onset), *n* (%) | 34 (27.0) |  |  |
| Young onset (≤ 40 years at onset), *n* (%) | 8 (6.4) |  |  |
| Duration of PD (range), *months* | 6 – 168 |  |  |
| Duration of PD (mean ± SD), *months* | 57.2 ± 35.3 |  |  |
| Duration of PD (median), *months* | 48 |  |  |
| Family history of tremor or parkinsonism, *n* (%) | 20 (15.9) | 0 (0%) |  |
| Hoehn and Yahr stage at study (median) | 2.0 |  |  |
| Hoehn and Yahr stage at study (range) | 1 – 5 |  |  |

### LRRK2 G2019S mutation analysis in PD cases and controls

The LRRK2 p.G2019S mutation was absent in all subjects. All 126 PD patients and 54 controls were homozygous for the p.G2019S (GG) normal allele. Allelic frequencies for p.G2019S (rs34637584) SNP were in Hardy-Weinberg equilibrium (P = 1). KASP assay performed very well. 100% call rates was achieved with clear genotyping clusters and no ambiguous calls.

## DISCUSSION

The data from this second cohort of PD from Nigeria corroborate the conclusions from our earlier publication that the commonest pathogenic mutation in LRRK2, (itself the commonest genetic cause of both familial and sporadic PD) the p.G2019S mutation, observed in several different populations, is not a frequent cause of PD in Nigeria. (7) This is similar to the view from other studies in the black populations of sub Saharan Africa, and in contrast to the data from Northern African Arab and Berber populations. In a small cohort study from Zambia (east Africa), the p.G2019S mutation was absent in 39 PD patients (10), as was the observation in another group of 54 PD patients and 46 controls from Ghana (west Africa).(11) In a study of 205 South African PD patients from varied backgrounds (42% Caucasian, 31% Afrikaner, 17% mixed ancestry, 8% Black and 2% Indian), LRRK2 p.G2019S mutation was not found in Afrikaner and Black South African patients. Only 2% of patients (three Caucasian and one patient from a mixed ancestry) were found to be carriers (12). All patients shared a common ancestor with the other haplotype 1-associated families reported worldwide.(12) This ethnic and geographical disparity is not peculiar to the African continent, and has been reported worldwide. (13) For example, Papapetropoulos *et al* explored the frequency of the G2019S LRRK2 mutation in southern Europe to determine if it was uniformly common, and found no patients with the mutation in mainland Greece, despite the close proximity to other European regions with high prevalence of LRRK2 mutations. (14) The disparity has been attributed to the LRRK2 p.G2019S gene mutation being a result mutational events in a common ancestral founder, at least one of which occurred in the Near East about 4000 years ago.(15-17) The genetic architecture of black sub Saharan Africans is evidently different from that of other populations within Africa (particularly North Africa), and also on other continents.

Our findings, together with those of other researchers from sub Saharan Africa lead us to reiterate the strong need to continue to explore the genetic underpinnings of apparently sporadic and inherited PD in persons of black African ancestry. This will necessitate incorporating larger cohorts to ensure that studies are sufficiently powered to reveal any significant genetic traits when contrasted with otherwise healthy controls. Furthermore, such studies should preferably employ advanced high throughput genetic technologies (such as whole genome and whole exome sequencing) to identify SNPs and other molecular targets associated with PD, and improve the likelihood that the complex underpinnings of the genetic component of PD in sub Saharan Africa are unravelled. This is particularly important to ensure that down stream applications of the results of genomics studies, such as genetic test batteries developed for PD in the future, will have improved yield, and genetic counselling will be developed with content relevant to the sub Saharan African population. In addition, such data will provide a basis for expanding our understanding of the genetics mechanisms contributing to PD in our population.

Our study utilized a rigorous clinical and laboratory methodology for case ascertainment and genotyping respectively. Diagnosis of PD was based on the MDS criteria, and all cases also fulfilled the UKPD Brain Bank criteria, verified in person by two neurologists specializing in movement disorders. With respect to the genotyping, the KASP is considered to be a reliable and more cost-effective genotyping assay for p.G2019S screening than multiplex methods, with the added advantage of being time-effective.(18)

We conclude that the contribution of genetics to the causation of PD in sub Saharan Africa remain unclear and largely unexplored. The region offers the unique advantage of having the greatest diversity of polymorphisms than any other world region, and this can be explored through adequately funded collaborative studies.

